# Pyramiding panicle-level heat avoidance and grain-level heat tolerance improves rice grain appearance under high-temperature grain filling

**DOI:** 10.64898/2026.08.20.745907

**Authors:** Hirofumi Fukuda, Toshihiro Sakamoto, Jun-ichi Yonemaru, Daisuke Ogawa

## Abstract

High temperature during grain filling increases rice grain chalkiness and deteriorates grain appearance under climate warming. Although several loci that reduce chalkiness have been identified, breeding strategies that integrate grain-level heat tolerance with panicle-level heat avoidance remain limited. Here we characterized SL2033, a chromosome segment substitution line carrying a long IR64-derived segment on chromosome 10, and evaluated the combination of the chromosome 10 segment with *Appearance quality of brown rice 1* (*Apq1*), a quantitative trait locus associated with reduced heat-induced chalkiness that acts at the grain level. Compared with its recurrent parent Koshihikari, SL2033 had longer flag leaves, altered vertical plant architecture, and lower panicle temperature. Total starch and protein contents were comparable between the two genotypes, whereas RNA-seq analysis of the developing endosperm identified specific differences in heat-, stress-, and cell wall–related transcripts. In a two-year field trial, a pyramided line combining the SL2033-derived segment with *Apq1* had the highest proportion of perfect grains and lowest frequencies of multiple chalky-kernel types during the year with hotter grain-filling conditions, with no detectable yield penalty. The pyramided line combined longer flag leaves, as in SL2033, with shorter panicle exsertion, as in an *Apq1* near-isogenic line, and had the lowest panicle temperature among the tested genotypes. Time-series unmanned aerial vehicle imaging also detected genotype-dependent differences in plant height during early grain filling, supporting distinct temporal patterns of plant development among the lines. These findings demonstrate that pyramiding genetic loci that confer panicle-level and grain-level heat tolerance is a promising strategy for improving rice grain appearance under high-temperature field conditions, which are becoming increasingly prevalent.

## 1. Introduction

Rice grain filling is highly vulnerable to heat stress, which can reduce productivity and deteriorate grain appearance. During grain filling, heat increases the formation of chalky kernels, including milky-white, white-belly-and-white-back, and basal-white grains, thereby decreasing the proportion of visually perfect grains (Shimoyanagi et al., 2021; Xu et al., 2021). The occurrence of chalky kernels increases markedly at the average air temperature reaches 27 °C or higher during the first 20 days after heading (DAH) (Wakamatsu et al., 2007; reviewed by Morita et al., 2016). The precise timing of heat exposure is critical. The grain-filling stage between 4 and 16 DAH is particularly sensitive to heat stress, although the exact timing differs among chalkiness types. Milky-white kernels are most readily induced by heat exposure at around 12 DAH, whereas the induction of white-back kernels peaks at around 16 DAH (Tashiro and Wardlaw, 1991). Heat stress during the early grain-filling stage disrupts endosperm development and water distribution, resulting in chalky grain formation (Ishimaru et al., 2009). Because grain appearance strongly influences the market value and acceptance of rice cultivars, reducing the occurrence of heat-induced chalkiness is an important breeding target under increasingly warm field conditions (Murata et al., 2014; Ishimaru et al., 2022; Fukuda et al., 2025; reviewed by Zhao et al., 2022).

The heat load experienced by developing grains is determined by ambient air temperature and the panicle temperature. Panicle temperatures can differ from air temperature depending on radiation, humidity, wind, transpiration, and plant architectural traits (Yoshimoto et al., 2011; Fukai and Mitchell, 2022; Ishimaru et al., 2022). Consequently, rice with different genotypes grown under the same air temperature may experience different levels of reproductive-stage heat load. Panicle temperature together with plant architecture and environmental measurements is a useful indicator of the thermal conditions experienced by developing grains (Fukai and Mitchell, 2022; Ishimaru et al., 2022; Tian et al., 2024).

Chromosome segment substitution lines (CSSLs) are useful genetic resources for dissecting quantitative traits related to heat resilience because donor genomic regions can be evaluated in a common genetic background (Ali et al., 2010; Nagata et al., 2023). In our previous study, three CSSLs showed reduced chalkiness and improved grain appearance compared with their recurrent parent, the *japonica* cultivar Koshihikari (Fukuda et al., 2025): among these, SL2033, which carries a 1×10^—6^–13.45 Mb chromosomal segment derived from the *indica* cultivar IR64 on chromosome 10 and short ones on several other chromosome, had the highest percentage of perfect grains during the hottest year of a 3-year field trial (Fukuda et al., 2025). However, the causal factor within this introgressed region remains unidentified. It is also unclear whether the superior grain appearance of SL2033 is associated with differences in storage-compound accumulation, panicle heat exposure, plant architecture, or early endosperm transcriptional responses relevant to subsequent chalkiness formation (Yamakawa et al., 2007; Liu et al., 2010; Zhao et al., 2024).

A separate question related to breeding is whether the IR64-derived chromosomal segment in SL2033 (hereafter referred to as the SL2033-derived segment) can provide additional value when combined with a previously characterized locus for grain appearance under heat stress. *Appearance quality of brown rice 1* (*Apq1*) was identified from the *indica* cultivar Habataki as a QTL allele on chromosome 7 associated with an increased percentage of perfect grains under heat stress in the Koshihikari genetic background (Murata et al., 2014). Fine-mapping and functional analysis identified the gene for *Sucrose synthase 3* (*Sus3*) as the causal gene underlying *Apq1*, with the Habataki allele showing enhanced expression under heat stress and contributing to improved grain appearance (Takehara et al., 2018). *Apq1/Sus3* is one of the best-characterized genetic factors associated with reduced heat-induced chalkiness in rice (reviewed by Zhao et al., 2022), but it remains unknown whether combining *Apq1* with the SL2033-derived segment can further improve grain appearance under field conditions. Evaluating this combination would determine whether the advantage conferred by SL2033 can complement the established effect of *Apq1*.

In this study, we aimed to assess the phenotypic associations and breeding value of the SL2033-derived segment and its combination with *Apq1*. We characterized the grain composition, panicle temperature, plant architecture, and early endosperm transcript profiles associated with the reduced chalkiness of SL2033. Next, we evaluated a pyramided line derived from SL2033 and an *Apq1* near-isogenic line (NIL) to determine whether their combination could further improve grain appearance under field conditions.

## 2. Materials and methods

### 2.1. Resequencing analysis of SL2033

Using the CTAB method (Murray and Thompson, 1980), we extracted genomic DNA from leaf blades of SL2033 (Nagata et al. 2015), which harbors chromosomal segments derived from the *indica* cultivar IR64 and had superior grain appearance in our previous study (Fukuda et al., 2025). Whole-genome resequencing reads from SL2033 were quality-filtered and trimmed using Trimmomatic v0.39 (Bolger et al., 2014), and the cleaned paired-end reads were aligned to the Nipponbare rice reference genome assembly IRGSP-1.0 using BWA-MEM (Li 2013) with default parameters. SAM files were converted to BAM format, merged, sorted, and indexed using SAMtools (Li et al., 2009). Duplicates were marked using Picard, and local realignment around indels was subsequently performed using the Genome Analysis Toolkit (GATK; McKenna et al., 2010). Variants were called from the realigned BAM files using GATK v4.0.12.0. Heterozygous genotype calls and calls with a read depth (DP) below 5 were flagged using ‘--genotype-filter-expression “isHet == 1”‘ and ‘--genotype-filter-expression “DP < 5”‘, respectively, and excluded from subsequent analyses. Indels were removed, and SNPs were identified separately for Koshihikari and IR64 relative to Nipponbare (IRGSP-1.0). The two SNP datasets were then compared to identifying biallelic loci that differentiated Koshihikari from IR64. 104,259 high-confidence SNP loci distinguishing the two parental genotypes were retained and used to identify Koshihikari- and IR64-derived genomic regions and to construct the graphical genotype of SL2033. The parental origin of each chromosomal segment was inferred from the SNP genotypes and plotted according to marker positions across the 12 rice chromosomes. Continuous regions with IR64-type SNPs were defined as donor-derived segments, and their physical boundaries were estimated from the nearest flanking SNP markers.

### 2.2. Rice plant materials and cultivation conditions

#### 2.2.1. Development of the SL2033 × *Apq1*-NIL pyramided line

During development of the pyramided line, rice plants were cultivated in a biotron breeding system (Tanaka et al., 2016). Unpollinated spikelets of *Apq1*-NIL (Takehara et al., 2018) were emasculated by incubation in a water bath at 43 °C for 7 min. The upper half of the hull was then cut with scissors. Pollen from bright yellow dehiscent anthers of SL2033 was used to directly pollinate the emasculated spikelets. To prevent further pollination, panicles bearing the pollinated spikelets were enclosed in envelopes. Mature F_1_ seeds were collected 30–40 days after pollination. The F_2_ population was obtained from eight F_1_ plants. To develop a pyramided line carrying both the IR64-derived region of SL2033 and the Habataki-derived *Apq1* region, 35 F_2_ lines were genotyped using codominant DNA markers, and plants homozygous for both target regions were selected. The *Apq1* region was genotyped with 5′-TTCAGCCAAGAACAGAACAGTGG-3′ (forward primer, Chr. 7: 25,473,624–25,473,646 in the Koshihikari genome) and 5′-CTTCTCTTCATCCTCCTCCTTGG-3′ (reverse primer, Chr. 7: 25,473,767–25,473,789), which amplified a 165-bp fragment. The SL2033-derived region was genotyped with 5′-CACATCATCTGACTAGATCCCATC-3′ (forward primer, Chr. 10: 10,429,090–10,429,113 in the Koshihikari genome) and 5′-TGAGTGCAAGAAACTTGTCATTT-3′ (reverse primer, Chr. 10: 10,429,209–10,429,231), which amplified a 141-bp fragment. One F_2_ line homozygous for both target regions was selected as the pyramided line.

#### 2.2.2 Rice cultivation in the field

Plants were grown in paddy fields in Kannondai, Tsukuba City, Ibaraki Prefecture, Japan, in 2024 (36°01’29.4"N, 140°06’28.4"E, 21 m a.s.l.) and 2025 (36°01’28.7"N, 140°06’37.3"E, 23 m a.s.l.). Seeds were soaked in water at 30 °C for 2 days, sown in trays (on April 22, 2024 and on April 21, 2025) filled with synthetic granular soil (Sumitomo Chemical Company Co., Ltd., Tokyo, Japan), and incubated at 30 °C in the dark for 2 days. Those trays were placed in a field for 1 month and then transplanted by hand (18 cm apart × 30 cm apart, planting density 18.5 hills/m^2^) in the field. Fertilizer was applied at 59:59:45 kg/ha (N:P_2_O_5_:K_2_O) in 2024 and at 56:176:56 kg/ha in 2025. Plants were grown from late-May to September. Data on daily average temperature during cultivation were obtained from the Japan Meteorological Agency. The mean air temperature during 0–20 DAH exceeded 27 °C in both years (Fig. S1).

### 2.3. Phenotypic analysis

#### 2.3.1. Manual assessment of agronomic traits

Agronomic traits were evaluated as described in Fukuda et al. (2025) and Fukuda et al. (2026a). For culm length, panicle length, panicle number, plant height, panicle exsertion, and flag leaf length and width, one biological replicate (*n*) was defined as a single plant. Length-and width-related traits were measured on the longest culm using a ruler from 5 DAH onward in at least 6 plants per genotype. Plant height was determined as the length of the main culm from the soil surface to the tip of the panicle. Panicle exsertion was defined as the distance from the flag leaf lamina joint to the panicle neck node. Panicle number was counted after heading was complete and before harvest. For aboveground dry weight, stem and leaf dry weight, panicle dry weight, and brown rice weight, *n* was defined as a bulk sample of 6 plants. Three to six biological replicates per genotype were used for aboveground, stem and leaf, and panicle dry-weight measurements, and four for brown rice weight per plant. Mature shoots were air-dried in a greenhouse, their aboveground dry weight was measured, and the shoots were cut 3 cm below the panicle base to separate panicles from stems and leaves. Panicles were weighed, and stem and leaf weight was calculated by subtracting panicle weight from aboveground dry weight. The bulked panicles were threshed, and brown rice weight per plant was calculated by dividing the total brown rice weight of each bulk sample by six (the number of plants in the sample).

#### 2.3.2. Assessment of grain appearance

Grain (brown rice) appearance was assessed from 96-dpi scanned images using a grain discriminator (RGQI 100B, Satake Corp., Hiroshima, Japan). Four biological replicates were analyzed per genotype, with at least 1900 grains analyzed per replicate. Following Fukuda et al. (2025), perfect grains were defined as non-chalky kernels with normal grain morphology (i.e., undamaged and fully mature grains). Chalky kernels were classified into the milky-white, basal-white, and white-belly-and-white-back types.

#### 2.3.3. Measurement of total starch and protein contents in brown rice

Four biological replicates were used per genotype, with each replicate consisting of six-plant bulk sample. Starch content was measured using an EnzyChrom Starch Assay Kit (E2ST-100; BioAssay Systems, Hayward, CA, USA), according to the manufacturer’s protocol. Brown rice grains (≥100 per replicate) were ground into a fine powder. For each biological replicate, starch was extracted from approximately 10 mg of powder and measured in four technical replicates, and their average was used as the value for the biological replicate. Protein content in approximately 1000 grains from each six-plant bulk sample (one biological replicate) was measured with a near-infrared grain tester (AN-820; Kett Electric Laboratory Co., Ltd., Tokyo, Japan), according to the manufacturer’s protocol. Instrument performance was verified using a rice grain standard (protein content, 6.3%; moisture content, 13.5%; amylose content, 17.6%) provided by the manufacturer.

#### 2.3.4. UAV measurement of plant height

Plant height was estimated using RGB images acquired by a Phantom 4 Pro V2 unmanned aerial vehicle (UAV) (P4V2; DJI, Shenzhen, China), following a previously described structure-from-motion multi-view stereo workflow (Fukuda et al., 2026b). Images were acquired at approximately weekly intervals from before transplanting (ground surface after leveling) through the ripening stage (primary focus of this study) to the harvest stage and processed using Agisoft Metashape Professional v.2.2.0 (Agisoft, St. Petersburg, Russia) to generate digital surface models (DSMs) for each acquisition date. For each plot, the ground surface height was defined as the mean DSM value obtained before transplanting. Plant height was then calculated as the 95th percentile of pixel-wise height difference between the DSMs at each acquisition date and the corresponding ground surface height within each 30 × 18-cm grid (corresponding to the footprint of an individual plant) along a transect.

#### 2.3.5. Thermal imaging of rice plants

The temperatures of panicles at 13–20 DAH were measured using photographs taken by an infrared thermal imaging camera (R300S; Nippon Avionics Co., Ltd., Yokohama, Japan) at around 9:30 am (air temperature at the time of imaging: 35.1 °C in 2024, 34.5 °C in 2025). The measurement wavelength was 8–14 μm and the spatial resolution was 1.2 mrad. The detectable temperature range was −40 to 120 °C, and the temperature resolution at 30 °C was 0.03 °C. The data were analyzed using a multifunctional reporting program, InfReC Analyzer NS9500 Standard (Nippon Avionics), and the maximum temperature of one panicle was used as one biological replicate.

### 2.4. Transcriptome analysis

#### 2.4.1. Endosperm sampling

For caryopsis sampling, spikelets in the field were marked on the day of flowering and collected at 4 days after flowering (DAF) (i.e., 5 DAH), which corresponded to approximately 15 weeks after sowing in 2024. The 4 DAF timepoint was selected because this stage corresponds to early endosperm development, before extensive starch accumulation, when transcriptional responses associated with subsequent chalkiness formation can be detected (Yamakawa et al., 2007; Liu et al., 2010; Shi et al., 2024). At least 10 spikelets were taken from the upper thirds of three panicles collected from different plants from 9:00 to 10:00 am, immediately frozen in liquid nitrogen, combined into one biological replicate, and stored at −80 °C. The husk was removed from frozen caryopses. Endosperms were separated from the embryonic bud on the cold base and then stored at −80 °C.

#### 2.4.2. RNA extraction and RNA-seq analysis

Frozen samples were crushed in a 3D bead-type homogenizer (ShakeMaster® AUTO, BMS-A20TP; Bio Medical Science Inc., Tokyo, Japan), and total RNA was immediately extracted from each sample using an RNeasy Plant Mini Kit (Qiagen, Hilden, Germany), according to the manufacturer’s protocol.

RNA-sequencing (RNA-seq) library preparation, sequencing, read preprocessing, mapping, read counting, and analysis of differentially expressed genes (DEGs) were performed as described in Fukuda et al. (2026a). Briefly, mRNA-seq libraries were prepared using the NEBNext Ultra II Directional mRNA-seq kit for Illumina (New England Biolabs, Ipswich, MA, USA) and sequenced on an Illumina NovaSeq X Plus platform with paired-end 150-bp reads. Reads were trimmed and mapped to the rice reference genome “Os-Nipponbare-Reference-IRGSP-1.0” using CLC Genomics Workbench version 25.0 (Qiagen), and uniquely mapped exon reads were counted. Expressed genes were defined as those with raw count ≥1 in each of the library samples. Identification of DEGs was based on raw read counts and was performed using the R package DESeq2 (version 1.42.1) in R software (version 4.3.2) as genes with |log_2_(FC)| > 1 and adjusted *P* value (padj) < 0.05 vs. the control group. Regularized log-transformed counts generated using the rlog function in DESeq2 were used for principal component (PC) analysis. The first and second PCs (PC1 and PC2) were visualized using the plotPCA function in DESeq2.

#### 2.4.3. Gene ontology enrichment analysis

Gene ontology (GO) analysis of DEGs was conducted using ShinyGO v0.85 (Ge et al., 2020) with ‘osativa_eg_gene’ as the database ID and ‘Oryza sativa Japonica Group genes IRGSP-1.0’ as the genome assembly. Upregulated and downregulated DEGs were analyzed separately. Enriched GO terms with an FDR-adjusted *P* value <0.05 were retained and ranked by fold enrichment.

## 3. Results

### 3.1. Meteorological conditions during grain filling and characterization of SL2033

Mean air temperature during 0–20 DAH of Koshihikari was 28.6 °C in 2024 and 28.0 °C in 2025; that during the critical period of 4–12 DAH (i.e., early grain-filling) was 28.6 °C in 2024 and 27.5 °C in 2025 (Fig. S1).

The CSSL SL2033 carried several IR64-derived chromosomal segments in the Koshihikari genetic background; the largest segment was on chromosome 10 (Fig. 1A). Representative images of brown rice harvested in 2024 showed visually apparent chalky kernels in Koshihikari, particularly milky-white and white-back grains, whereas these defects were reduced in SL2033 (Fig. 1B), consistent with our previous findings (Fukuda et al., 2025). Neither total starch nor total protein content of brown rice differed significantly between SL2033 and Koshihikari under the tested conditions (Table S1). The maximum panicle temperature of SL2033 was significantly lower than that of Koshihikari at both 13 and 20 DAH (Fig. 2A, B). Plant height and flag leaf length were significantly higher and culm length–to– plant height ratio was significantly lower in SL2033 (Fig. 2C). Panicle exsertion and flag leaf width did not differ significantly between SL2033 and Koshihikari.

**Fig. 1.**
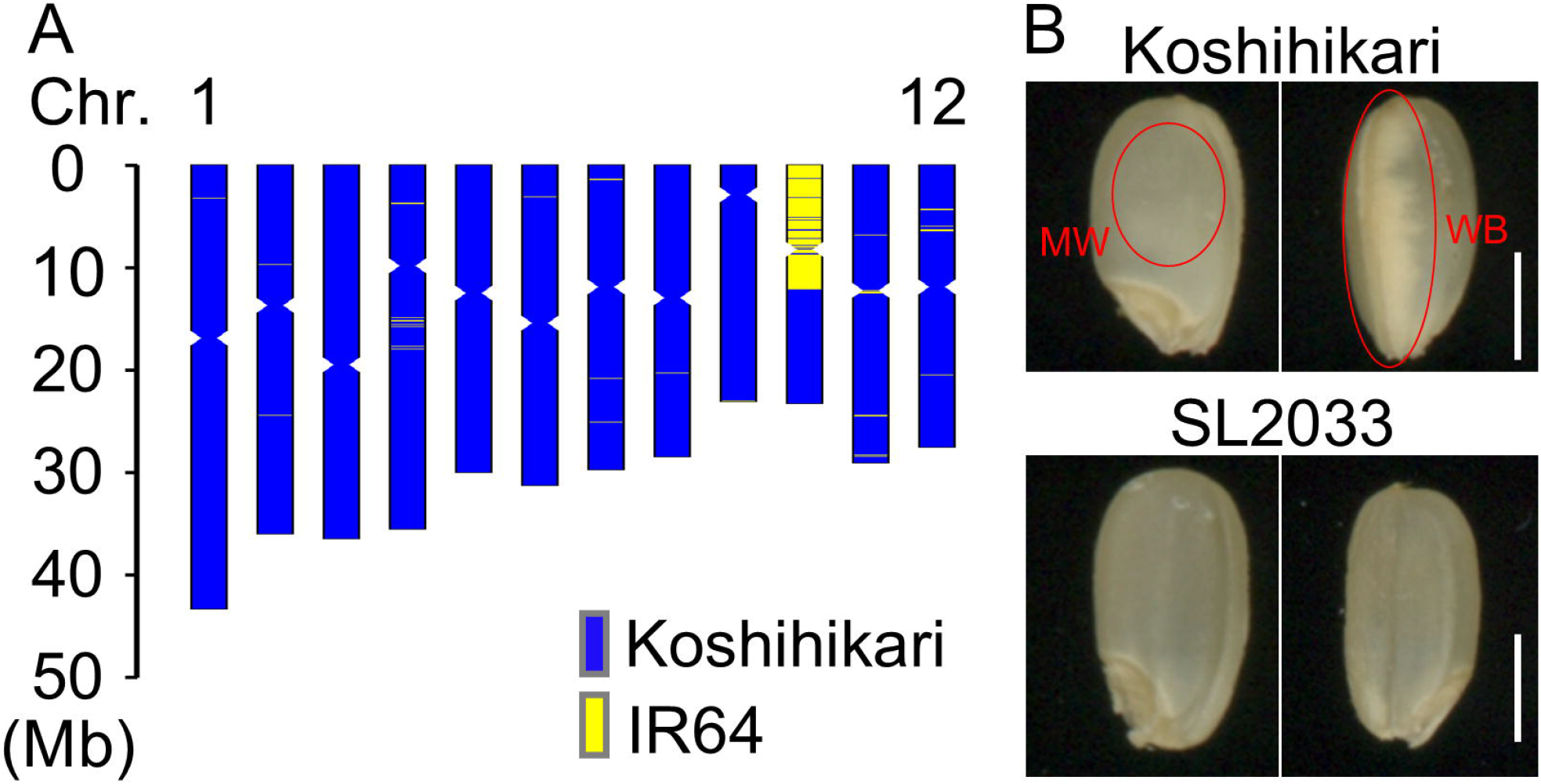
Graphical genotype of chromosome substitution line SL2033 and its grain appearance. (A) Graphical genotype. Genomic regions from *japonica* Koshihikari are in blue and *indica* IR64 are in yellow. (B) Appearance of brown rice grains of SL2033 and Koshihikari. SL2033 showed superior grain appearance, with reduced frequencies of milky-white (MW) and white-back (WB) kernels compared with Koshihikari. Scale bars indicate 2 mm. **basal-white (BW)**,

**Fig. 2.**
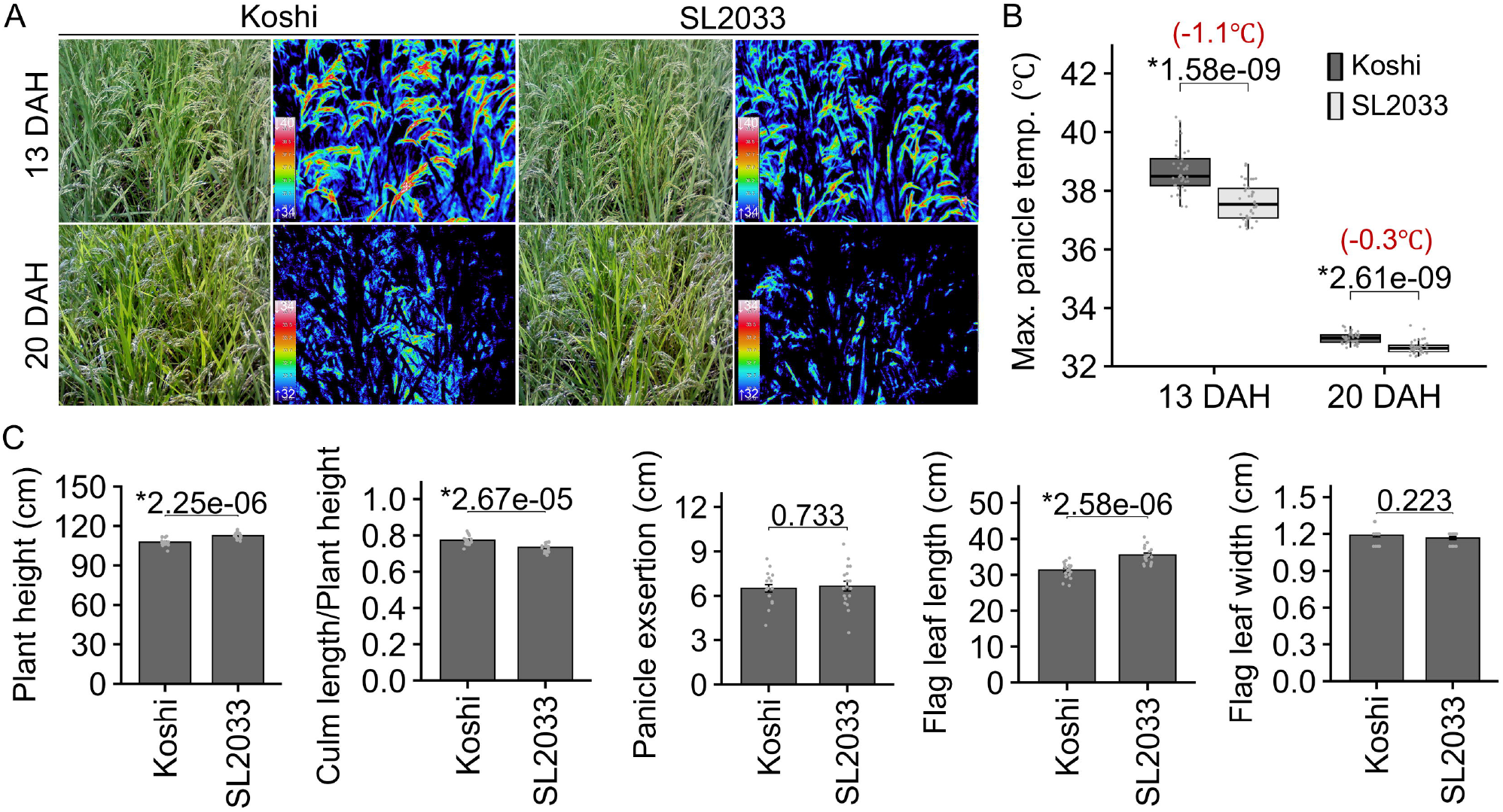
Plant morphology and panicle temperature in Koshihikari (Koshi) and SL2033. (A) Panicle temperature detection at 13 and 20 days after heading (DAH) in 2024. These photographs were taken at air temperature of 35.1 °C. Thermal images are displayed using pseudo-color scales ranging from 32–34°C at the lower end to 40°C at the upper end; blue/green colors indicate lower temperatures, whereas yellow/red/white colors indicate progressively higher temperatures. (B) Maximum panicle temperature at 13 and 20 DAH. Boxes indicate the interquartile range (25th–75th percentiles), horizontal lines within boxes indicate medians, whiskers indicate 1.5 × the interquartile range, and gray points represent individual observations. Temperature values in red are differences in mean values between Koshi and SL2033. (C) Plant height, culm length–to–plant height ratio, panicle exsertion, flag leaf length, and flag leaf width. Data are mean values ± SD (panicle temperature, *n* = 40; others, *n* = 18). *P* values (Student’s *t*-test) are shown for each comparison; asterisks indicate significant differences between Koshi and SL2033 (*P* < 0.05).

### 3.2. Altered endosperm transcript profiles in SL2033 at early grain filling

To examine the genome-wide effects of the IR64-derived chromosomal segment(s) on transcriptomes at early grain filling, we performed RNA-seq analysis of the endosperm at 4 DAF in SL2033 and Koshihikari. In total, 25,551 genes were expressed. In PC analysis, the endosperm transcriptome profiles visibly differed between SL2033 and Koshihikari (Fig. 3A). We identified 35 upregulated and 95 downregulated genes in SL2033 relative to Koshihikari (Fig. 3B). In GO enrichment analysis, upregulated genes were enriched in stress-related categories, including responses to reactive oxygen species, hydrogen peroxide, and inorganic substances, whereas downregulated genes were enriched in categories related to hydrolase activity acting on glycosyl bonds including cellulase activity (Fig. 3C). Two HSP20/alpha-crystallin family genes, *HSP23*.*2* (Os04g0445100/LOC_Os04g36750) and *HSP21*.*9* (Os11g0244200/LOC_Os11g13980) (Nyasulu et al., 2025), were included in the enriched GO term “response to reactive oxygen species” (GO:0000302) and were upregulated in SL2033 (Fig. 3D). Two cellulase genes, *CEL9A* (Os01g0220100/LOC_Os01g12070) and *GH9B7* (Os02g0151300/LOC_Os02g05744) (Yoshida et al., 2006; Xie et al., 2013), were included in the enriched GO term “cellulase activity” (GO:0008810) and were downregulated in SL2033 (Fig. 3E). Transcript levels of more than 10 heat shock factor genes were lower in SL2033 than in Koshihikari, although the magnitude and statistical support varied among the genes (Fig. 3F). These results identified genotype-dependent differences in early endosperm transcript profiles, including selective changes in stress- and cell wall–related transcripts.

**Fig. 3.**
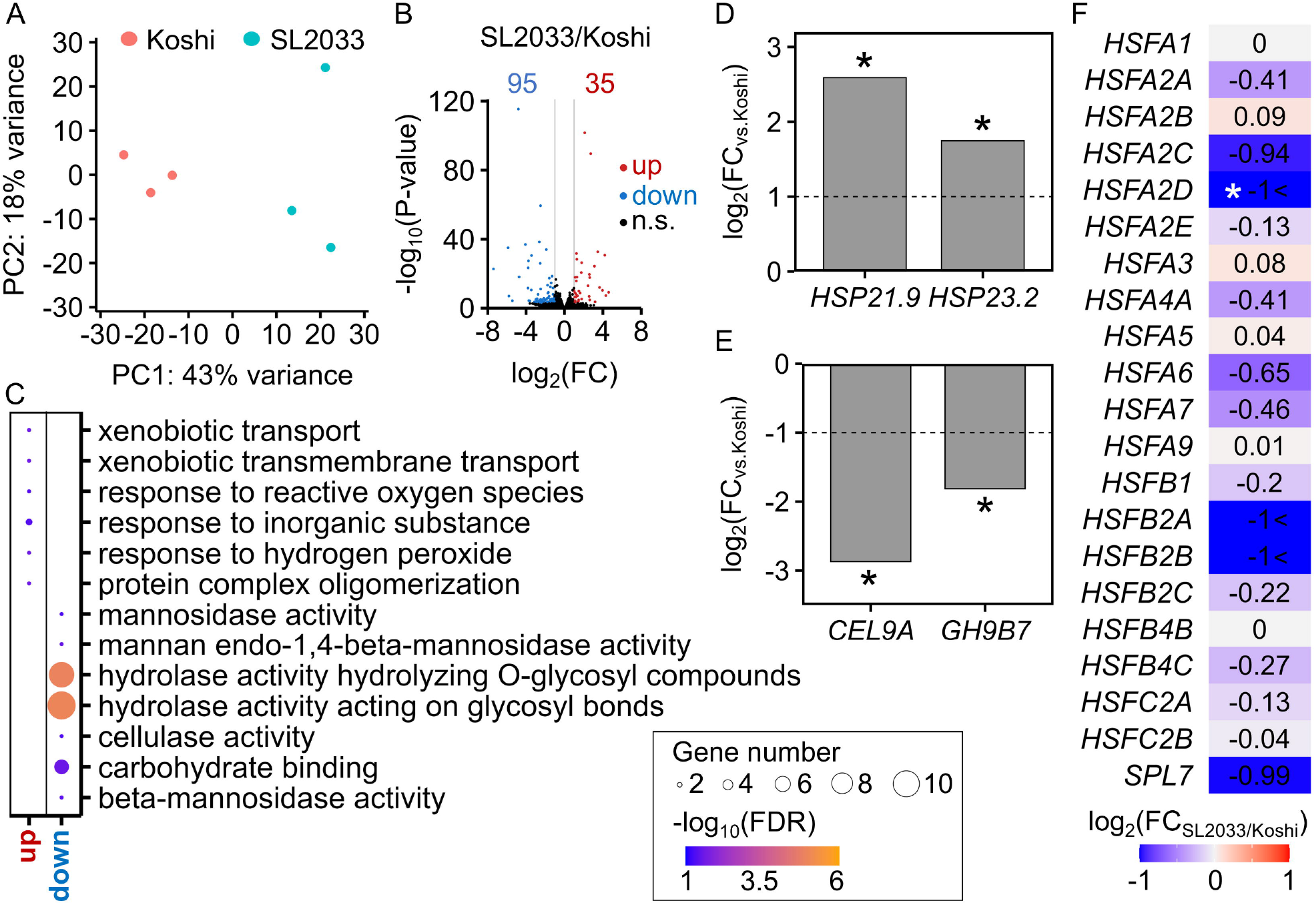
Endosperm transcript profiles of Koshihikari (Koshi) and SL2033. (A) Principal component (PC) analysis of endosperm transcriptomes at 4 days after flowering (i.e., 5 days after heading) in 2024. Individual data for PC1 and PC2 are plotted (*n* = 3). (B) Volcano plot of upregulated (red) and downregulated (in blue) genes at 4 days after flowering (5 days after heading) in SL2033 relative to Koshi. Genes whose RNA-seq counts were significantly different [|log_2_(fold change, FC)_SL2033/Koshi_)| > 1, padj *<* 0.05, *N* = 25,551] were counted. Log_2_(FC) was calculated using mean RNA-seq counts of three biological replicates for all the detected genes (raw read count ≥ 1 in each of the library samples). (C) Analysis of gene ontology (GO) biological process enrichment for genes significantly upregulated and downregulated in SL2033 relative to Koshi. Ten top-ranked GO terms per gene category are shown. The color bar describes enrichment −log_10_(FDR) values. (D–F) Changes in mRNA levels of selected genes related to heat and cell wall responses: (D) heat shock protein genes, (E) cellulase genes, and (F) heat shock factor genes. Asterisks indicate statistically significant differences in SL2033 relative to Koshi (|log_2_(FC)| > 1 and padj < 0.05).

### 3.3. Grain appearance in the SL2033 × *Apq1* pyramided line

The percentage of perfect grains and frequency of chalky kernels were higher in Koshihikari than in SL2033, Apq1-NIL, and SL2033 × *Apq1*-NIL, whereas the pyramided line had the highest percentage of perfect grains and lowest frequencies of the three chalky-kernel types in 2024, the hotter of the two years (Fig. 4A, B; Fig. S1); a similar but less pronounced trend was observed in 2025 (Table S2). Grain width was comparable among the lines, whereas grain length was higher and grain thickness was lower in the pyramided line than in Koshihikari (Table S3). No significant difference in brown rice weight per plant was detected (Fig. 4C, indicating no detectable yield penalty.

**Fig. 4.**
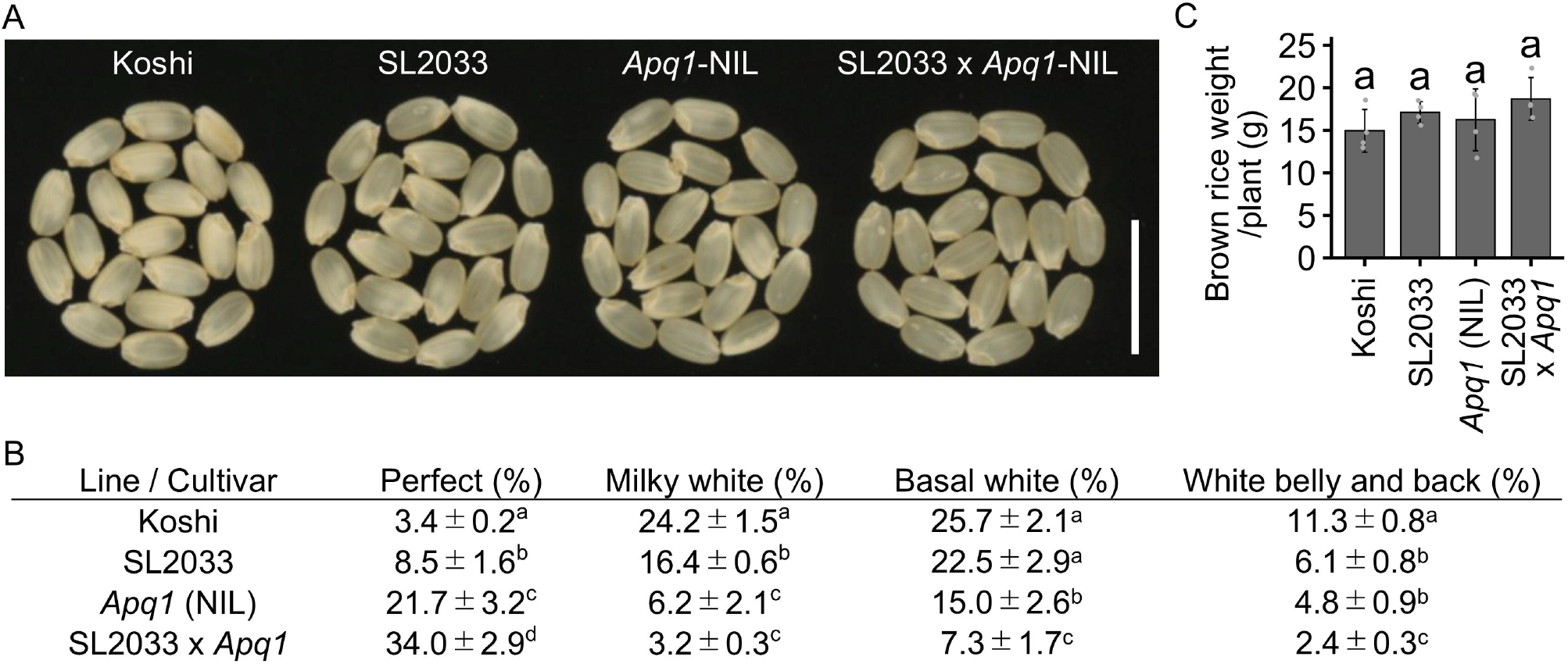
Appearance of brown rice. (A) Grains of rice plants in 2024. Koshihikari (Koshi) grains were the most chalky and the least transparent. Scale bar, 10 mm. (B) Percentages of perfect and chalky kernels. (C) Brown rice weight per plant at harvest in Koshi, SL2033, *Apq1*-NIL, and the pyramided line (SL2033 × *Apq1*-NIL). Individual data points are plotted. In (B) and (C), the means ± SD are shown (*n* = 4). Different lowercase letters indicate significant differences among each line and Koshi (*P* < 0.05, Tukey–Kramer test).

### 3.4. Panicle temperature and architectural traits in the SL2033 × *Apq1* pyramided line

Panicle temperature during grain filling was lowest in the pyramided line and highest in Koshihikari among the four genotypes (Fig. 5A, B). In time-series UAV measurements, SL2033 was significantly taller than Koshihikari from 1 to 8 DAH (this difference remained detectable at several later points), *Apq1*-NIL was significantly shorter than Koshihikari from 1 DAH, and the height of the pyramided line did not differ significantly from that of Koshihikari from 8 DAH to 22 DAH (Fig. 5C, Fig S2). In manual measurements, the pyramided line had longer flag leaves, as in SL2033, and shorter panicle exsertion, as in *Apq1*-NIL (Fig. 5D). No consistent changes in other agronomic traits were observed in the pyramided line across the two years (Fig. S3). These results indicate that the pyramided line had reduced panicle temperature during grain filling and integrated contrasting parental architectural traits without changes in flag leaf width.

**Fig. 5.**
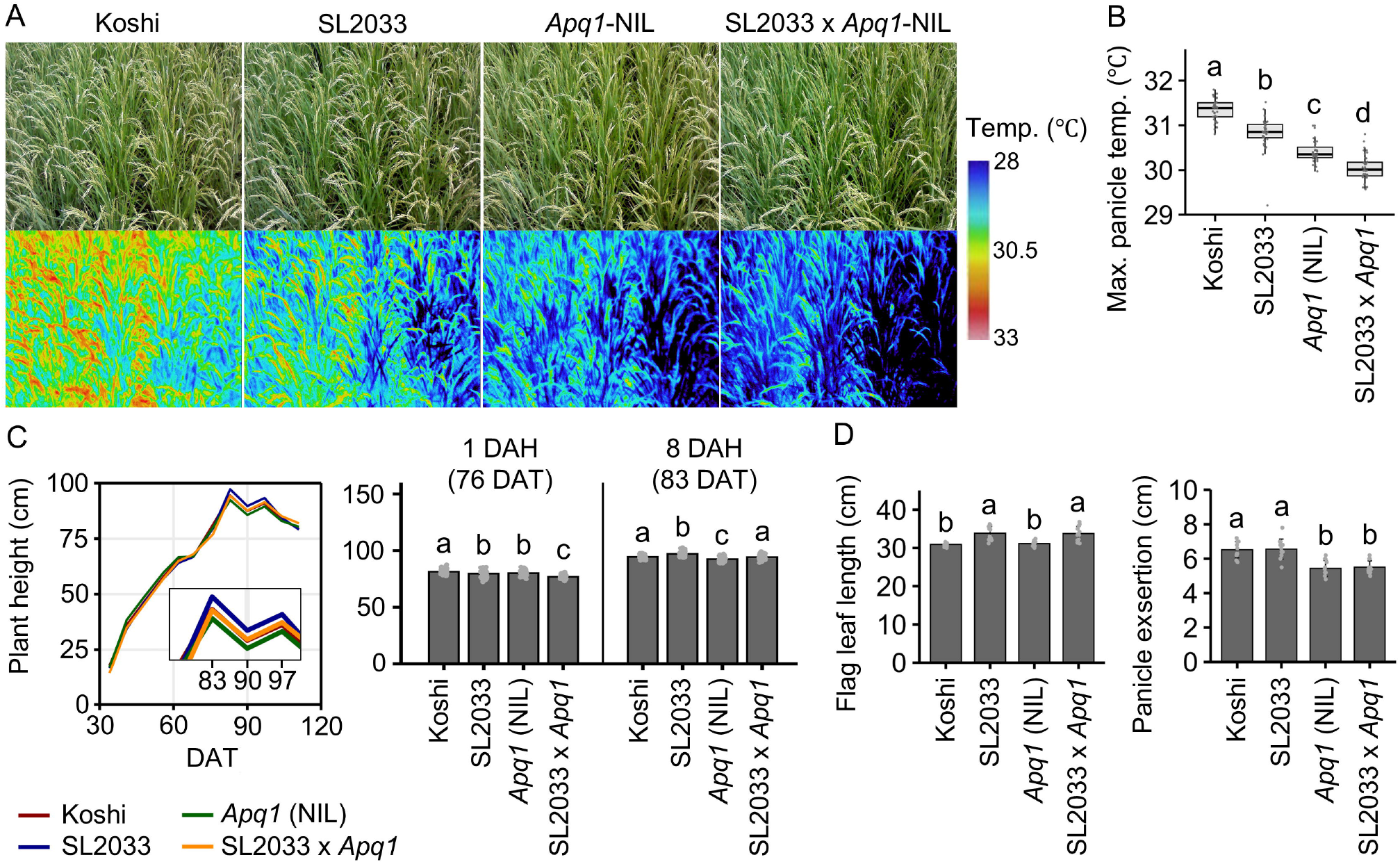
Panicle temperature and plant morphology in Koshihikari (Koshi) and lines with reduced chalkiness. (A) Panicle temperature at around 20 days after heading (DAH) in 2025; air temperature was 34.5 °C. (B) Maximum panicle temperature at around 20 DAH. (C) Plant height determined by UAV measurements at different days after transplanting (DAT); DAH values are indicated for Koshihikari. (D) Flag leaf length and panicle exsertion. Data are mean ± SD (panicle temperature, *n* = 40; plant height, *n* = 63; others, *n* = 12). Different lowercase letters indicate significant differences among all lines (*P* < 0.05, Tukey–Kramer test).

## 4. Discussion

### 4.1. Plant architecture–associated heat avoidance in SL2033

Despite no significant differences in total starch or protein content of brown rice, SL2033 had superior grain appearance (Figs. 1B, 4B, Table S1), indicating that the reduced chalkiness of SL2033 was not attributable to major changes in the bulk accumulation of storage compounds. Chalkiness is generally associated with heat-induced disruption of endosperm development rather than simply with changes in the total accumulation of storage compounds such as starch or protein (Kaneko et al., 2016; Wada et al., 2019). Consistent with this interpretation, the lower panicle temperature observed in SL2033 at grain filling (Fig. 2A, B) indicates that developing grains experienced a reduced heat load under field conditions. Longer flag leaves and altered vertical organization (Figs. 2C, 5C) may influence radiation interception, airflow, and the positional relationship between leaves and panicles, thereby contributing to a cooler panicle microenvironment. Panicle temperature is affected by canopy microclimate and heat-balance processes; panicles positioned lower within the canopy can remain cooler because of reduced solar radiation interception (Yoshimoto et al., 2011; Kitajima et al., 2022); our data are consistent with this interpretation. Likewise, in rice cultivar Nijinokirameki, canopy structure and leaf traits have been proposed to reduce panicle temperature during grain filling (Ishimaru et al., 2022). Thus, the superior grain appearance of SL2033 may be partly associated with plant architecture–mediated heat avoidance.

### 4.2. Endosperm transcriptional responses associated with reduced chalkiness in SL2033

Mitigation of heat-related disruption during endosperm development may reduce chalkiness (Kaneko et al., 2016; Wada et al., 2019). RNA-seq analysis at 4 DAF (during early grain filling) identified genotype-dependent differences in endosperm transcript abundance (Fig. 3). *HSP23*.*2* and *HSP21*.*9* had higher transcript levels in SL2033 than in Koshihikari. Because HSP20 proteins are associated with cell protection under heat stress (Nyasulu et al., 2025), these expression patterns may indicate differences in stress-related responses between SL2033 and Koshihikari. In contrast, the mRNA levels of several heat shock factor genes, which are key regulators of heat-responsive transcription and are typically induced by heat stress (Mittal et al., 2009), were lower in SL2033 than in Koshihikari (Fig. 3F). These contrasting patterns suggest selectivity in the changes in particular stress-related transcripts in SL2033. *CEL9A* and *GH9B7* were among the downregulated genes assigned to cellulase activity categories (Fig 3C, E). They are annotated as GH9/endoglucanase-related genes. As plant endoglucanases are involved in cell wall modification and restructuring (Yoshida et al., 2006; Xie et al., 2013; Glass et al., 2015), their reduced mRNA levels may be associated with differences in cell wall–related processes or tissue remodeling in the developing endosperm of SL2033.

### 4.3. Complementary effects of SL2033- and *Apq1*-associated traits on grain appearance

We demonstrated that the long SL2033-derived chromosomal segment on chromosome 10 can be combined with *Apq1* to further improve grain appearance under field conditions. The mean temperature at 0–20 DAH exceeded the approximately 27 °C threshold associated with increased grain chalkiness (Wakamatsu et al., 2007; reviewed by Morita et al., 2016) in both years (Fig. S1), with the early grain-filling period being hotter in 2024 than in 2025. Under the hottest post-heading conditions, the pyramided line showed the highest percentage of perfect grains and the lowest frequencies of chalkiness among the tested lines (Fig. 4A, B; Fig. S1) without a significant reduction in brown rice weight per plant or other yield-related traits (Fig. 4C, Fig. S3), suggesting that the improved grain appearance was not simply caused by reduced grain production or a smaller sink load. The favorable performance of the pyramided line may partly reflect the combination of distinct architectural traits associated with SL2033 and *Apq1*-NIL: SL2033 had longer flag leaves and greater plant height, whereas *Apq1*-NIL had shorter plant height in the UAV time-series analysis, and shorter panicle exsertion was observed as an *Apq1*-NIL-associated trait in the pyramided line (Figs. 2C; 5C, D). Panicle temperature is influenced by radiation interception, transpiration, airflow, and the positional relationship between panicles and surrounding leaves (Yoshimoto et al., 2011; Fukai and Mitchell, 2022; Ishimaru et al., 2022; Kitajima et al., 2022; Tian et al., 2024). Therefore, the combination of longer flag leaves and reduced panicle exsertion in the pyramided line may have altered the panicle microenvironment during grain filling. Consistent with this interpretation, the pyramided line had the lowest panicle temperature among the four genotypes (Fig. 5A, B). These results strongly suggest that SL2033- and *Apq1*-NIL-associated architectural traits can be integrated within a single genetic background and may jointly contribute to reduced panicle heat exposure. The breeding value of this combination may also extend beyond canopy-level heat avoidance. Under high temperature, Habataki-derived *Apq1*, corresponding to the highly expressed allele of *Sus3*, improves grain appearance through effects on sucrose synthase activity and carbohydrate metabolism (Murata et al., 2014; Takehara et al., 2018). *Sus3* has also been implicated in sucrose utilization and partitioning between conducting tissues and developing grains (Shi et al., 2024). These established functions provide a plausible explanation of the contribution of *Apq1* to the superior grain appearance of the pyramided line. The regulatory relationship and potential pathway interactions between the SL2033-derived leaf architecture and *Apq1* should be investigated in future studies.

### 4.4. Integrating panicle-level heat avoidance and grain-level heat tolerance for improved grain appearance

Most genetic studies of chalkiness have identified and selected loci on the basis of visible grain phenotypes, such as the percentage or area of chalky endosperm (Guo et al., 2011; Peng et al., 2014; Yang et al., 2022). Yang et al. (2024) pyramided two to four low-chalkiness QTLs and showed that chalkiness progressively decreased as favorable loci accumulated; whether these effects involved panicle-level heat avoidance or grain-level heat responses remained unresolved. Previous studies of *Apq1/Sus3* established its role in improving grain appearance through heat-responsive sucrose synthase activity during ripening but did not examine panicle temperature, panicle exsertion, or other heat-avoidance-related architectural traits (Murata et al., 2014; Takehara et al., 2018; Shi et al., 2024). The present study associated an introgression of *Apq1* with shorter panicle exsertion and reduced panicle temperature, and combined *Apq1* with a genetic resource associated with distinct heat avoidance–related architecture. The novel approach we used here was to extend conventional chalkiness-QTL pyramiding toward the integration of heat resilience traits at both the panicle microenvironment and developing-grain levels.

## Statements

### Data availability statement

The datasets used and/or analyzed in the current study are available from the corresponding author upon reasonable request.

### Authorship contribution

**Hirofumi Fukuda:** Writing – original draft, Writing – review & editing, Investigation, Data curation, Formal analysis, Visualization, Conceptualization. **Toshihiro Sakamoto:** Writing – review & editing, Investigation, Data curation. **Jun-ichi Yonemaru:** Writing – review & editing, Data curation, Formal analysis. **Daisuke Ogawa:** Writing – review & editing, Investigation, Data curation, Funding acquisition, Supervision.

## Acknowledgements

We thank Dr. Kazumasa Murata of the Toyama Prefectural Agricultural, Forestry and Fisheries Research Center for providing *Apq1*-NIL seeds. We thank Di Guan, Makiko Suzuki, Yuko Aono, and Aimi Murakami for their technical and field support. We appreciate the technical staff of the Institute of Crop Science, NARO, for their help in managing the rice fields. We thank Akari Fukuda for sharing the protocol of RNA-seq library preparation. We also thank Kett Electric Laboratory Co. Ltd. for helping with the analysis of protein content of rice grains. We are thankful to the editors at ELSS, Inc. (https://elss.co.jp/en/) for their professional editing services before submission.

## Declaration of generative AI and AI-assisted technologies in the writing process

During the preparation of this manuscript, we used DeepL Write, Microsoft Copilot, and ChatGPT to improve the grammar, spelling, and readability. After using these tools, the authors carefully reviewed and edited the content as needed and take full responsibility for the content of the published article.

