## Supplementary Figures for "Pyramiding panicle-level heat avoidance and grain-level heat tolerance improves rice grain appearance under high-temperature grain filling"

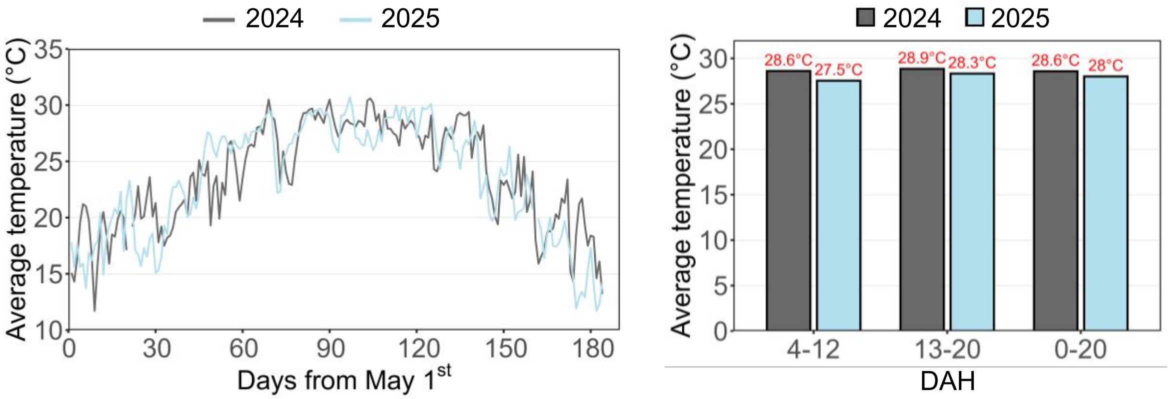


**Fig. S1 Daily temperature from May 1 to October 31 and average temperature at several days after heading (DAH) of Koshihikari in the field condition in 2024 and 2025.** The occurrence of chalky kernels increases when the average air temperature during the first 20 days after heading exceeds to 27° C (Wakamatsu et al., 2007; reviewed by Morita et al., 2016). DAH among lines used in this study was at most 3-d differences in both 2024 and 2025.

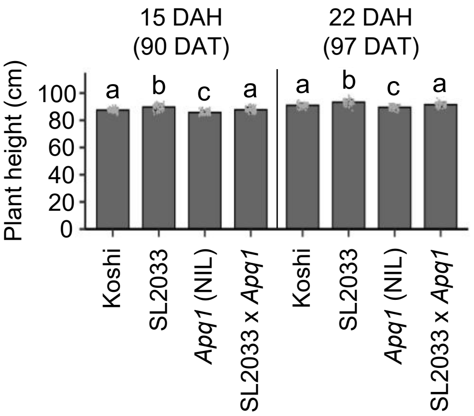


**Fig. S2. Plant height determined by UAV measurements at 15 and 22 days after heading (DAH).** DAH values are indicated for Koshihikari in 2025. Data are mean ± SD (*n* = 63). Different lowercase letters indicate significant differences among lines (*P* < 0.05, Tukey–Kramer test). DAT, days after transplanting.

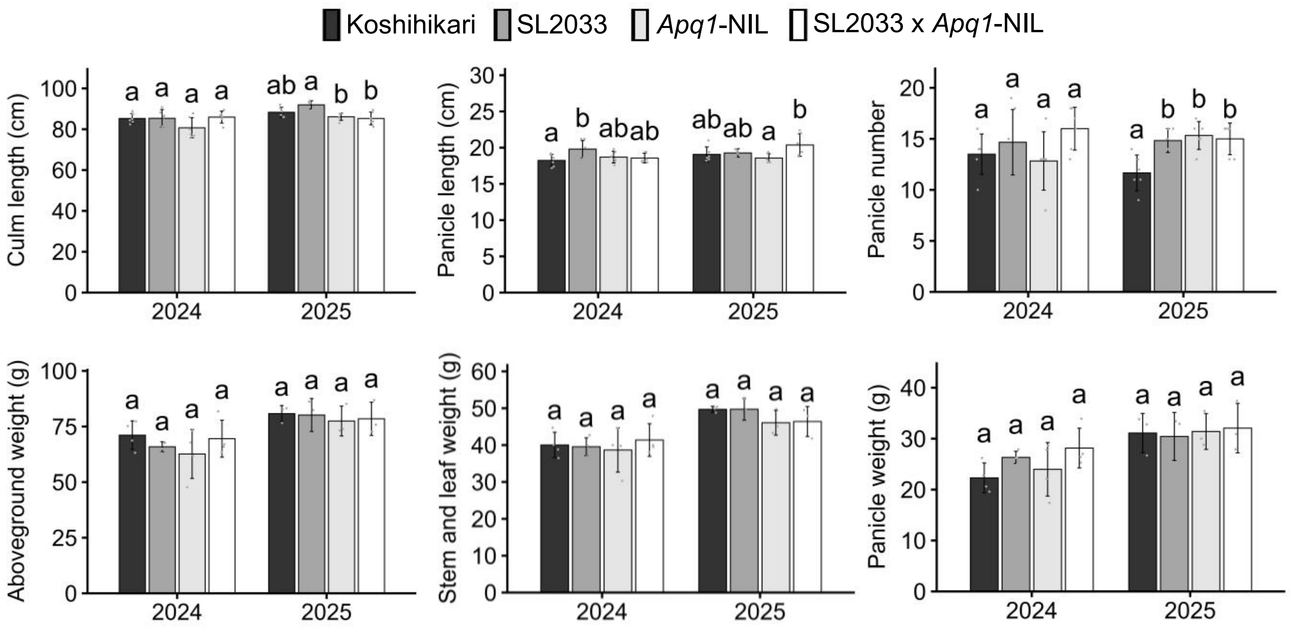


**Fig. S3. Other agronomical traits of each line and cultivar in 2025.** Culm length, panicle length, panicle number, aboveground (dry) weight, stem and leaf (dry) weight, and panicle (dry) weight at/after harvesting. Bar plots mean values ± SD (*n* = 3–6). Different letters above each bar indicate significant differences among each line and a Koshihikari genetic background (*P* < 0.05, Tukey-Kramer test).
